# Immunity conferred by drug-cured experimental *Trypanosoma cruzi* infections is long-lasting and cross-strain protective

**DOI:** 10.1101/741462

**Authors:** Gurdip Singh Mann, Amanda F. Francisco, Shiromani Jayawardhana, Martin C. Taylor, Michael D. Lewis, Francisco Olmo, Elisangela Oliveira de Freitas, Fabiana M. S. Leoratti, Cesar López-Camacho, Arturo Reyes-Sandoval, John M. Kelly

**Affiliations:** Department of Infection Biology, London School of Hygiene and Tropical Medicine, Keppel Street, London WC1E 7HT, UK; The Jenner Institute, Old Road Campus Research Building, Roosevelt Drive, Oxford OX3 7DQ, UK

## Abstract

**Background:** The long term and complex nature of Chagas disease in humans has restricted studies on vaccine feasibility. Animal models also have limitations due to technical difficulties in monitoring the extremely low parasite burden that is characteristic of chronic stage infections. Advances in imaging technology offer alternative approaches that circumvent these problems. Here, we describe the use of highly sensitive whole body *in vivo* imaging to assess the efficacy of recombinant viral vector vaccines and benznidazole-cured infections to protect mice from challenge with *Trypanosoma cruzi*.

**Methodology/Principal Findings:** Mice were infected with *T. cruzi* strains modified to express a red-shifted luciferase reporter. Using bioluminescence imaging, we assessed the degree of immunity to re-infection conferred after benznidazole-cure. Mice infected for 14 days or more, prior to the initiation of treatment, were highly protected from challenge with both homologous and heterologous strains (>99% reduction in parasite burden). Sterile protection against homologous challenge was frequently observed. This level of protection was considerably greater than that achieved with recombinant vaccines. It was also independent of the route of infection or size of the challenge inoculum, and was long-lasting, with no significant diminution in immunity after almost a year. When the primary infection was benznidazole-treated after 4 days (before completion of the first cycle of intracellular infection), the degree of protection was much reduced, an outcome associated with a minimal *T. cruzi*-specific IFN-γ^+^ T cell response.

**Conclusions/Significance:** Our findings suggest that a protective Chagas disease vaccine must have the ability to eliminate parasites before they reach organs/tissues, such as the GI tract, where once established, they become largely refractory to the induced immune response.

**AUTHOR SUMMARY:** Chagas disease, which is caused by the protozoan parasite *Trypanosoma cruzi*, is a major public health problem throughout Latin America. Attempts to develop a vaccine have been hampered by technical difficulties in monitoring the extremely low parasite burden during the life-long chronic stage of infection. To circumvent these issues, we used highly sensitive bioluminescence imaging to assess the ability of recombinant viral vector vaccines and drug-cured infections to confer protection against experimental challenge in mice. We observed that drug-cured infections were much more effective than subunit vaccines, with many instances of sterile protection. Efficacy was independent of the route of infection or size of the challenge inoculum, and was undiminished after almost a year. In addition, drug-cured infections conferred a high level of cross-strain protection. The highly sensitive imaging procedures enabled us to visualise parasite distribution in mice where sterile protection was not achieved. This suggested that to confer sterile protection, vaccines must prevent the infection of organs/tissues that act as parasite reservoirs during the chronic stage. Once established at these sites, parasites become largely refractory to vaccine-induced elimination.

## INTRODUCTION

Chagas disease is caused by the insect-transmitted protozoan *Trypanosoma cruzi* and is the most serious parasitic infection in the Americas. More than 5 million people are infected with this obligate intracellular parasite (1, 2), resulting in a financial burden estimated at $7 billion annually (3). In humans, the disease is characterised by an acute stage that occurs 2-8 weeks post-infection, during which bloodstream parasites are often detectable. Symptoms during this period are normally mild, although lethal outcomes can occur in 5% of diagnosed cases. The parasite numbers are then controlled by a vigorous adaptive immune response. However, sterile immunity is not achieved and infected individuals transition to a chronic stage, which in most cases, appears to be life-long (4). Around 30-40% of those infected eventually develop chronic disease pathology, a process that can take decades to become symptomatic. Cardiomyopathy is the most common clinical manifestation (5, 6), although 10-15% of people can develop digestive tract megasyndromes, sometimes in addition to cardiac disease.

Attempts to control Chagas disease have been challenging. For example, although public health measures have been successful in reducing disease transmission in several regions of South America, there is a vast zoonotic reservoir that complicates disease eradication by this route (7–9). The only drugs currently available to treat the infection, the nitroheterocycles benznidazole and nifurtimox, have limited efficacy and cause toxic side effects that can impact on patient compliance (10, 11). There have been no new treatments for almost 50 years, but progress in discovering new chemotherapeutic agents is now being accelerated by a range of drug development consortia encompassing both the academic and commercial sectors (12). For many years, vaccine development against Chagas disease has been inhibited by concerns that autoimmunity could play a role in disease pathogenesis (13, 14). Although not excluded as a contributory factor, the current consensus is that the risk has been overstated, and that the continued presence of the parasite is required to drive disease pathology (15–17).

The host response to *T. cruzi* infection involves a complex combination of both innate and adaptive immune mechanisms (18, 19). The innate system is key to controlling parasite proliferation and dissemination during the initial stages of infection (20), with important roles for both Toll-like receptor (TLR)-mediated inflammatory responses and TLR-independent processes (21). As the acute phase progresses, the development of an antigen-specific immune response, in which CD8^+^ IFN-γ^+^ T cells are the key effectors (18, 22), is the critical step in controlling the infection. In both humans and mice, the major targets of this cellular response are a small set of immunodominant epitopes within specific members of the trans-sialidase super-family of surface antigens (23). The observation that the pattern of this recognition displays strain variation has been interpreted as indicative that immune evasion could be operating at a population level. The adaptive response reduces the parasite burden by >99%, with the infection becoming highly focal, and in BALB/c mice at least, confined predominantly to the large intestine, stomach, and to a lesser extent, the gut mesentery tissue and sites in the skin (24, 25). The reason why the immune system is not able to eradicate the infection is unresolved. It does not appear to involve exhaustion of the CD8^+^ IFN-γ^+^ T cell response, which continues to suppress, but not eliminate, the parasite burden throughout the long chronic stage (26). These findings have questioned the feasibility of developing an effective anti-T. *cruzi* vaccine.

Experimental vaccination of animal models against *T. cruzi* infection has a long history (27), although there have been few instances in which unequivocal sterile protection has been reported. Approaches have included the use of attenuated parasites (28, 29), immunisation with cell fractions (30), purified or recombinant proteins (31, 32) and the use of DNA vaccines. In the latter case, viral backbones based on vaccinia (33), yellow fever (34) and adenovirus (35) have been used to facilitate expression of a range of parasite antigens such as trans-sialidase, Tc24, and the amastigote surface protein-2 (ASP-2) (also a member of the trans-sialidase super-family). Reported outcomes include protection from lethal infection, reduction in the acute stage parasite burden, induction of a favourable cytokine profile, and reduction in disease pathology. However, detailed analysis of vaccine efficacy has been limited by an inability to accurately monitor parasite levels during the chronic stage and technical difficulties in assessing the effect of the immune response on tissue distribution following challenge infections.

Recently, *in vivo* imaging approaches have been exploited to provide new insights into infection dynamics during experimental chronic Chagas disease (24, 25, 36). These studies have revealed the pantropic nature of acute stage infections, and shown that during the chronic stage, the adaptive immune response restricts parasites to small infection foci, predominantly within the GI tract. Other tissues and organs, including the heart and skeletal muscle, are infected sporadically, the extent of which is influenced by host:parasite genetics and immune status. Additional factors such as nutrition, environmental stimuli, age and co-infections could also play a role in this complex chronic infection profile (37). The survival of the small parasite foci within apparently tolerant sites is crucial for long-term infection, although the immunological context of these reservoirs is unknown. Another contributor to the long-term nature of *T. cruzi* infections could be the phenomenon of parasite dormancy; individual intracellular amastigotes can enter an apparently quiescent state in which they cease to replicate and exhibit reduced drug sensitivity (38). Neither the mechanisms involved, nor the potential implications for immune evasion have yet been established.

Highly sensitive bioluminescence imaging involves the use of *T. cruzi* strains that have been modified to express a red-shifted luciferase reporter (39). The system allows the real-time monitoring of parasite burden in experimental mice during chronic stage infections. There is a robust correlation between parasite numbers and whole animal bioluminescence, with a limit of detection close to 100 parasites (24). Here, we describe the use of this imaging technology to assess the extent of protection in benznidazole-cured mice following re-challenge with homologous and heterologous strains. Our findings have important implications for vaccine strategies.

## MATERIALS AND METHODS

### Generation of recombinant ASP-2/TS vaccines

The fusion gene encoding the ASP-2 and TS peptides (Figure 1A) was generated by linking sequences corresponding to the mouse Ig kappa chain signal peptide, ASP-2 amino acids 1-694 (GenBank accession no. U77951) and TS amino acids 1-624 (GenBank accession no. L38457). The furin 2A splice site linker was inserted between the trypanosome sequences to ensure the subsequent generation of two separate peptides from a single open reading frame (40). The *ASP-2/TS* fusion gene was cloned into the ChAdOx1 (41) and MVA (42, 43) viral-vectored vaccine platforms, and confirmed by sequencing prior to use in protection studies.

### Assessment of recombinant vaccine immunogenicity

Vaccines were prepared in PBS and administered intramuscularly into the left and right quadriceps muscles of mice. The ChAdOx1 vaccine was administered at 1×10^8^ infectious units per dose. With MVA:ASP-2/TS, each dose was equivalent to 1×10^6^ plaque forming units. ELISpots were carried out using either peripheral blood mononuclear cells (PBMCs) or splenocytes. Briefly, MAIP ELISpot plates (Millipore) were coated at 4°C overnight with anti-mouse IFN-γ mAb AN-18 (Mabtech), at 250 ng per well, and then blocked for 1 h with complete DMEM medium (10% foetal calf serum). Whole blood was sampled by venesection of the tail vein and PBMCs were isolated using histopaque 1083 (Sigma), and plated at 5×10^5^ cells per well with 20-mer specific peptides overlapping by 10 amino acids (10 μg ml^-1^) (Pepscan Presto). Splenocytes from naïve mice were plated 2.5×10^5^ per well. After 16 h incubation, cells were discarded and plates washed with PBS. 50 μl of biotinylated anti-mouse IFN-γ mAb RA-6A2 (1:1000 in PBS) was then added to each well and incubated for 2 h. After another washing step, streptavidin peroxidase (Sigma) was added and incubated at 37°C for 1 h. The plates were washed and developed with TMB substrate solution (Mabtech). When spots were visible, the reaction was stopped by washing the plate with water. Spots were analysed using an ELISpot reader, and the number of spot-forming cells/10^6^ PBMCs producing IFN-γ was calculated.

### Murine infections and bioluminescence imaging

Animal work was performed under UK Home Office project licence (PPL 70/8207) and approved by the LSHTM Animal Welfare and Ethical Review Board. Procedures were in accordance with the UK Animals (Scientific Procedures) Act 1986 (ASPA). BALB/c mice were purchased from Charles River (UK), and CB17 SCID mice were bred in-house. Animals were maintained under specific pathogen-free conditions in individually ventilated cages. They experienced a 12 h light/dark cycle, with access to food and water *ad libitum*. SCID mice were infected with 1×10^4^ bioluminescent bloodstream trypomastigotes (BTs) in 0.2 ml PBS via intraperitoneal (i.p.) injection (24, 25, 36). BALB/c female mice, aged 8-10 weeks, were infected i.p with 1×10^3^ BTs derived from SCID mouse blood. At experimental end-points, mice were sacrificed by exsanguination under terminal anaesthesia.

For *in vivo* imaging, mice were injected with 150 mg kg^-1^ d-luciferin i.p., then anaesthetized using 2.5% (v/v) gaseous isoflurane. They were placed in an IVIS Lumina II system (Caliper Life Science) 5-10 min after d-luciferin administration and images acquired using LivingImage 4.3. Exposure times varied from 30 s to 5 min, depending on signal intensity. After imaging, mice were revived and returned to cages. For *ex vivo* imaging, mice were injected with d-luciferin, and sacrificed by exsanguination under terminal anaesthesia 5 min later. They were then perfused via the heart with 10 ml 0.3 mg ml^-1^ d-luciferin in PBS. Organs and tissues were removed and transferred to a Petri dish in a standardized arrangement, soaked in 0.3 mg ml^-1^ d-luciferin in PBS, and imaged using maximum detection settings (5 min exposure, large binning). The remaining animal parts and carcass were checked for residual bioluminescent foci, also using maximum detection settings (24, 25). To estimate parasite burden in live mice, regions of interest were drawn using LivingImage v.4.3 to quantify bioluminescence as total flux (photons/second), summed from dorsal and ventral images. The detection threshold for *in vivo* imaging was determined using uninfected mice.

### Drug treatment and immunosuppression

Benznidazole was synthesized by Epichem Pty Ltd., Australia, and prepared at 10 mg ml^-1^ in an aqueous suspension vehicle containing 5% (v/v DMSO, 0.5% (w/v) hydroxypropyl methylcellulose, 0.5% (v/v) benzyl alcohol and 0.4% (v/v) Tween 80. It was administered by oral gavage. To detect any residual infection following treatment, mice were immunosuppressed with cyclophosphamide monohydrate (Sigma) in D-PBS (200 mg kg^-1^), administered by i.p. injection every 4 days, for 3 doses. Two weeks after the end of immunosuppression, mice that were bioluminescence negative by both *in vivo* and *ex vivo* imaging were designated as cured.

## RESULTS

### Monitoring ASP-2/TS vaccine efficacy using highly sensitive bioluminescence imaging

The *T. cruzi* amastigote surface protein 2 (ASP-2) and trypomastigote cell surface protein trans-sialidase (TS) have shown promise as vaccine candidates against *T. cruzi* infections (44–46). To further assess their efficacy, we used two replication-deficient recombinant vaccine platforms, a chimpanzee adenovirus (ChAdOx1) and a Modified Vaccinia Ankara virus (MVA), expressing ASP-2 and TS peptides from a single open reading frame (Fig 1A) (Materials and Methods) (40). We used a homologous prime-boost vaccination strategy in which BALB/c mice received intramuscular injections administered one week apart (Fig 1B) and confirmed immunogenicity of the vaccine delivery system using an *ex vivo* IFN-γ^+^ ELISpot. Three weeks after receiving either a prime only, or a prime-booster vaccination, splenocytes were plated on antibody coated ELISpot plates. When these cells were then stimulated with a peptide pool representing the entire ASP-2/TS sequence, there was a pronounced increase in the number of peptide specific IFN-γ^+^ splenocytes, particularly in those mice that had received the booster (Fig 1C).

**Fig 1.**
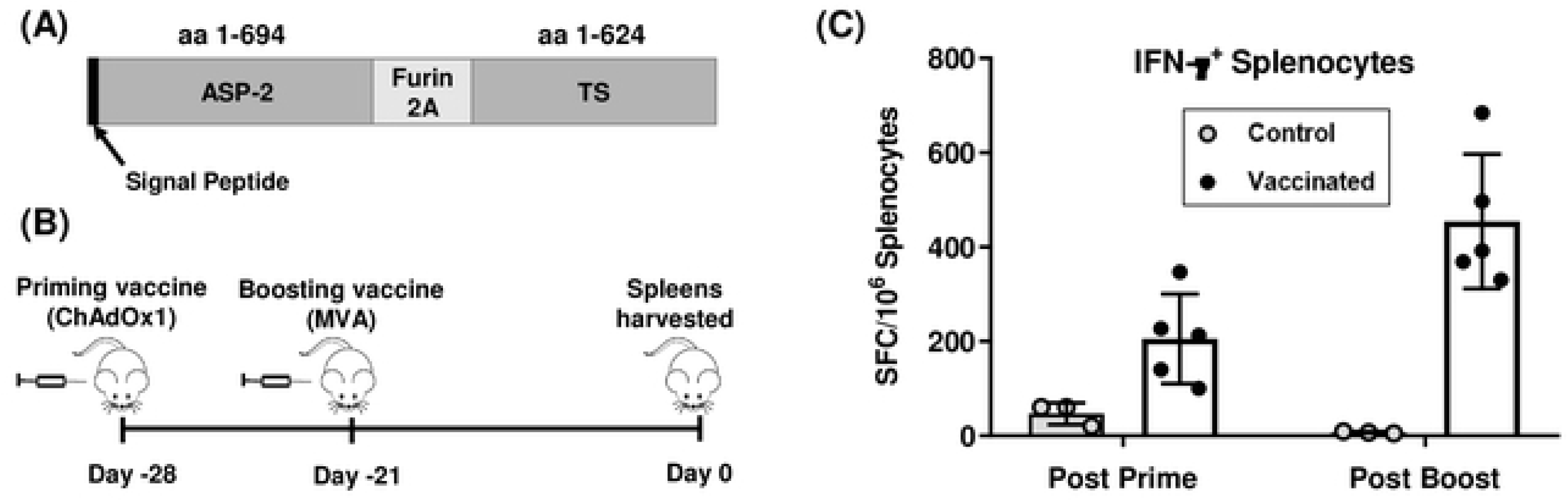
Immunogenicity of the recombinant ASP-2/TS vaccine. (A) Open reading frame encoding ASP-2 and TS peptides, with the mouse Ig kappa chain signal peptide and furin 2A splice site linker indicated. (B) Prime-boost vaccination strategy. BALB/c mice (n=5, per vaccinated group) were inoculated one week apart with ChAdOx1 (priming) and then MVA (boosting) recombinant vaccines containing the *ASP-2/TS* fusion gene. Control mice received vaccinations with the same viral vectors engineered to express dengue protein NS-1 (n=3). (C) ELISpot analysis. Splenocytes were plated onto antibody coated ELISpot plates and stimulated for 16 h with a peptide pool covering the entire ASP-2 and TS sequences (Materials and Methods). Data are presented as *ex vivo* IFN-γ spot forming cells (SFCs) per 10^6^ splenocytes.

To test protective efficacy, mice were vaccinated using the homologous prime-boost strategy outlined above. Three weeks after the MVA booster, they were challenged by i.p. injection with 10^3^ bioluminescent *T. cruzi* blood trypomastigotes (strain CL Brener) (Fig 2A). The resulting infection was monitored by *in vivo* imaging (36) (Materials and Methods). In BALB/c mice, parasites rapidly disseminate and proliferate, with the infection reaching a peak after approximately 2 weeks. Thereafter, a vigorous adaptive immune response reduces the parasite burden by >2 orders of magnitude, and the infection transitions to the life-long chronic stage (24). No differences were observed in the bioluminescence-inferred parasite burden between vaccinated and control mice at the earliest time-point assessed (day 7, post-infection) (Figs 2B and C). However, by day 15, the peak of the acute stage, the ASP-2/TS vaccinated mice displayed a 77% reduction in the bioluminescence-inferred parasite burden. This protective effect was maintained until day 21. By day 28, against a background of immune-mediated reduction in the parasite burden, there were no significant differences between the vaccinated and control groups. From that point onwards, the parasite burden remained similar between the groups (Figs 2B and C). Following termination of the experiment (day 95), *ex vivo* imaging of internal organs and tissues revealed that the profile of infection in the vaccinated cohort was typical of the chronic stage. The colon and/or stomach were the major tissues persistently parasitized, with infections in other organs being sporadic. There were no apparent differences in the tissue-specific parasite burden between vaccinated and control mice (Fig 2D). Therefore, although vaccination with the ASP-2/TS constructs can reduce parasite burden during the acute stage, it does not impact on the long term burden of chronic infections.

**Fig 2.**
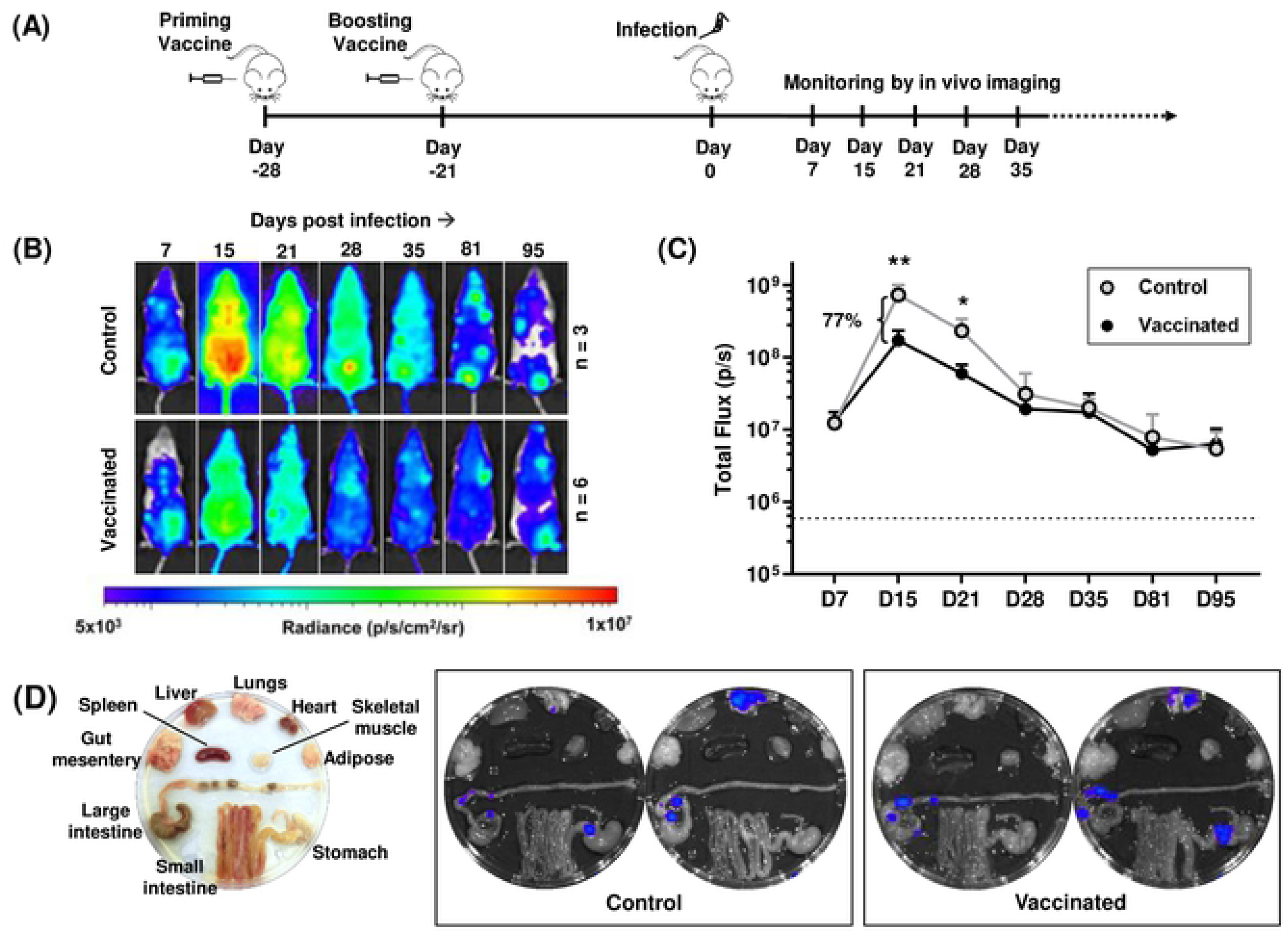
Assessing the efficacy of the ASP-2/TS vaccine to protect mice against *T. cruzi* infection. (A) Timeline of vaccination experiment. (B) Assessment of efficacy. Vaccinated (n=5) and control BALB/c mice (n=3) were infected i.p. with 10^3^ bioluminescent *T. cruzi* trypomastigotes and monitored by *in vivo* imaging. Representative ventral images (from a single mouse in each case) are shown at sequential time points post-infection. All images use the same log_10_-scale heat-map with minimum and maximum radiance values indicated. (C) Quantification of whole animal bioluminescence (ventral and dorsal) of vaccinated and control cohorts (mean + SD). Dashed line indicates background bioluminescence; (**) *p*<0.01, (*) *p*<0.05. (D) *Ex vivo* bioluminescence imaging. Left-hand image; arrangement of organs and tissues. Insets, representative images of organs from control and vaccinated mice, 95 days post-infection.

### Drug-cured T. cruzi infection confers significant protection against challenge with a homologous strain

To place the recombinant vaccine results into context, we sought to establish the extent to which drug-cured infections could enhance the capacity of the murine immune response to protect against challenge. BALB/C mice were first inoculated i.p. with bioluminescent parasites (CL Brener strain) (n=12). At three different points post-infection (4, 14 and 36 days) (Fig 3A), we initiated treatment with benznidazole (100 mg kg^-1^), once daily, for 20 consecutive days. This dosing regimen was shown to be curative (Supplementary Fig 1), in line with previous results (47, 48). The plasma concentration of benznidazole falls below the *in vitro* EC50 value in approximately 12 h (48). 21 days after the cessation of treatment, the cured mice were re-infected i.p. and monitored regularly by *in vivo* imaging (Fig 3B and C). 70 days after challenge, mice found to be bioluminescence-negative were immunosuppressed to facilitate the outgrowth and dissemination of any residual parasites, then assessed further by *ex vivo* imaging two weeks later (Fig 3D) (Materials and Methods).

**Fig 3.**
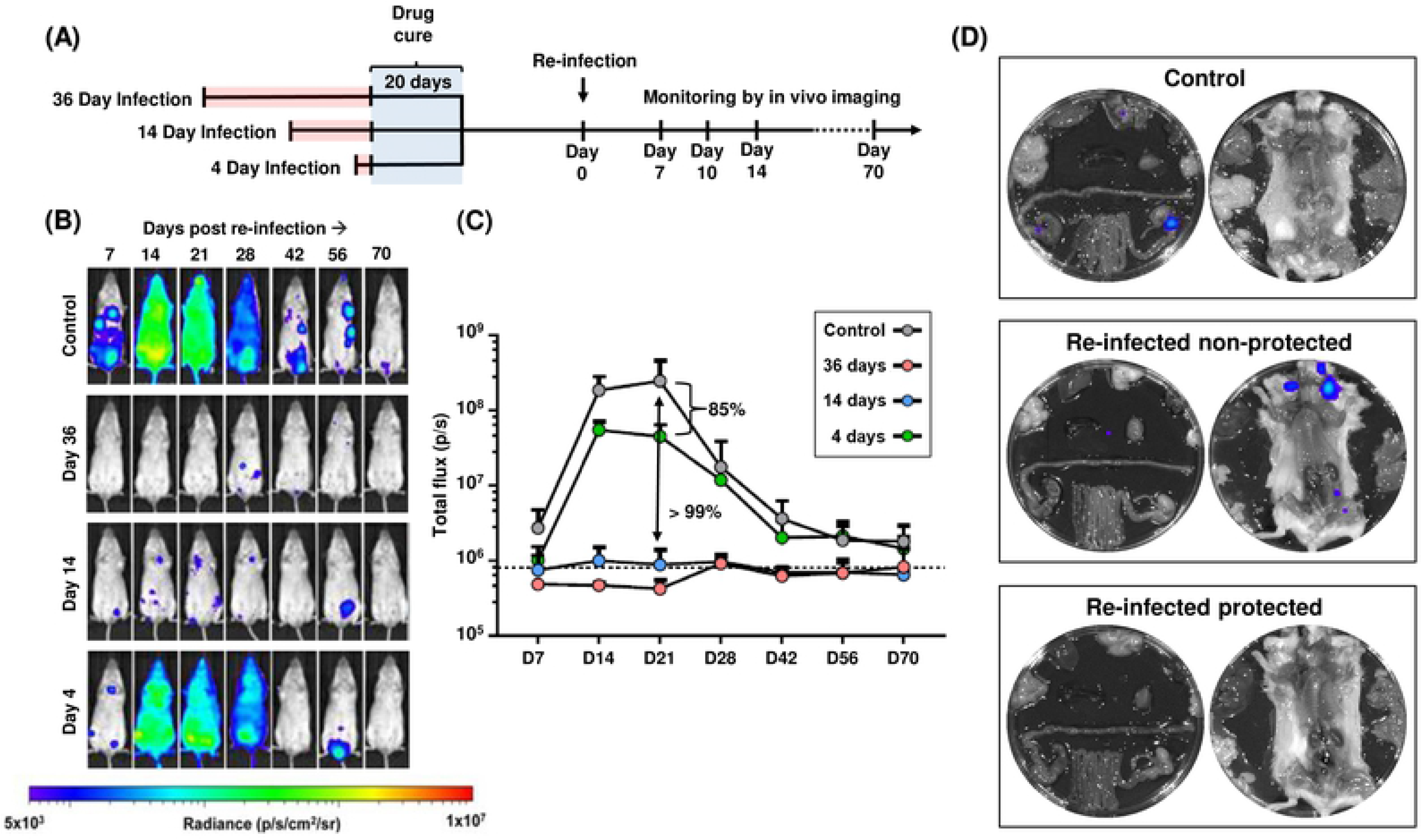
Drug-cured infections confer significant protection against challenge with a homologous *T. cruzi* strain. (A) Outline of strategy. BALB/c mice were infected i.p. with 10^3^ bioluminescent trypomastigotes (CL Brener strain) and subjected to curative benznidazole treatment initiated at various times post-infection. 21 days after the end of treatment, they were re-infected i.p. and monitored for a further 70 days. Bioluminescence-negative mice were then immunosuppressed using cyclophosphamide (Materials and Methods) and assessed by *ex vivo* imaging two weeks later. Results were derived from 2 independent experiments, each involving 6 mice per cohort. (B) Representative ventral images of benznidazole-cured mice following re-infection. The number of days at which drug treatment was initiated, following the primary infection, is indicated (left). All images use the same log10-scale heat-map with minimum and maximum radiance values indicated. (C) Total body bioluminescence (sum of ventral and dorsal images) of drug-cured mice following re-infection (n=12) (means ± SD) derived by *in vivo* imaging. The length of the primary infection is indicated (inset). (D) Representative *ex vivo* bioluminescence images. Upper inset, organs from control mouse 70 days post-infection. Central inset, organs from a mouse that was re-infected following curative treatment initiated on day 36 of the primary infection. On day 70 of the challenge infection, immunosuppressive treatment was initiated and the organs then harvested. In this instance, a residual infection was identified. Lower inset, organs from a mouse treated as above, which was non-infected after challenge.

When curative treatment was initiated at 36 days post-infection, none of the mice exhibited a distinct acute stage infection peak following challenge (Figs 3B and C). There was >99% reduction in the inferred parasite burden in all cases, compared to primary infection control mice. At the experimental end-point, 6 out of 12 mice were shown to be fully protected (Table 1, Fig 3D, Supplementary Fig 2). In cases where the primary infection was allowed to proceed for 14 days prior to the initiation of curative benznidazole treatment, the infection profile following challenge was very similar to the cohort where treatment was initiated 36 days post-infection. However, a greater number of foci were detectable during the period corresponding to the acute stage of primary infections (Fig 3B), with full protection achieved in 3 out of 12 mice. In contrast to both of the above, when curative treatment began 4 days after the primary infection, there was a clear acute stage peak in the bioluminescence profile following challenge (Fig 3C). The kinetic profile mirrored that in control infected mice, although the maximum parasite burden was 85% lower, and sterile protection was restricted to a single mouse (Table 1). Therefore, although the immune response induced by a short course infection is able to impact on the burden of re-infection, the effect is significantly limited compared with what is achievable when the primary infection is allowed to progress fully into the acute stage, prior to treatment.

**Table 1.** Summary of protection data

| Preliminary infection strain | Challenge strain | Route of infection | Challenge dose | Length of primary infection<sup>1</sup> | Period between cure and challenge | Mean parasite reduction<sup>2</sup> | Level of sterile protection |
| --- | --- | --- | --- | --- | --- | --- | --- |
| CL-Brener | CL-Brener | i.p. | 1 x 10<sup>3</sup> | 36 days | 21 days | >99% | 6/12<sup>3</sup> |
| CL-Brener | CL-Brener | i.p. | 1 x 10<sup>3</sup> | 14 days | 21 days | >99% | 3/12<sup>3</sup> |
| CL-Brener | CL-Brener | i.p. | 1 x 10<sup>3</sup> | 4 days | 21 days | 85% | 1/12<sup>3</sup> |
| CL-Brener | CL-Brener | s.c. | 1 x 10<sup>3</sup> | 36 days | 21 days | >99% | 4/5 |
| CL-Brener | CL-Brener | i.p. | 6 x 10<sup>4</sup> | 36 days | 21 days | >99% | 5/6 |
| CL-Brener | CL-Brener | i.p. | 1 x 10<sup>3</sup> | 36 days | 338 days | >99% | 4/6 |
| JR | JR | i.p. | 1 x 10<sup>3</sup> | 36 days | 20 days | >99% | 1/6 |
| JR | CL-Brener | i.p. | 1 x 10<sup>3</sup> | 36 days | 20 days | >99% | 0/6 |
| CL-Brener | JR | i.p. | 1 x 10<sup>3</sup> | 36 days | 20 days | >99% | 0/6 |

To assess whether this protective effect was dependent on the route of inoculation, we repeated the 36 day challenge experiment using the subcutaneous (s.c.) route. *T. cruzi* transmission normally occurs when parasite-infected faeces from the insect vector are rubbed into the wound produced by blood feeding; therefore, s.c. inoculation probably reflects more closely how the majority of human infections occur. The mice were inoculated s.c. with 10^3^ bloodstream trypomastigotes for both the primary and challenge infections, but the protocols and treatment timelines were otherwise identical to those used previously (Fig 3A, 36 day infection). In control mice, the bioluminescence profile of s.c. infections, and the resulting organ-specific tropism during the chronic stage was similar to that in i.p. infections (Figs 4A, B and C), as shown previously (24, 25). When the challenge cohort was assessed by *in vivo* imaging (Fig 4A), none of the mice displayed an acute stage peak, and bioluminescence was at, or close to background levels. At the experimental end-point, all of the mice were immunosuppressed and then subjected to post-mortem *ex vivo* organ imaging to test for foci of infection below the limit of *in vivo* detection. 4 out of 5 were found to be bioluminescence negative in all analyses and were designated as protected (Table 1).

**Fig 4.**
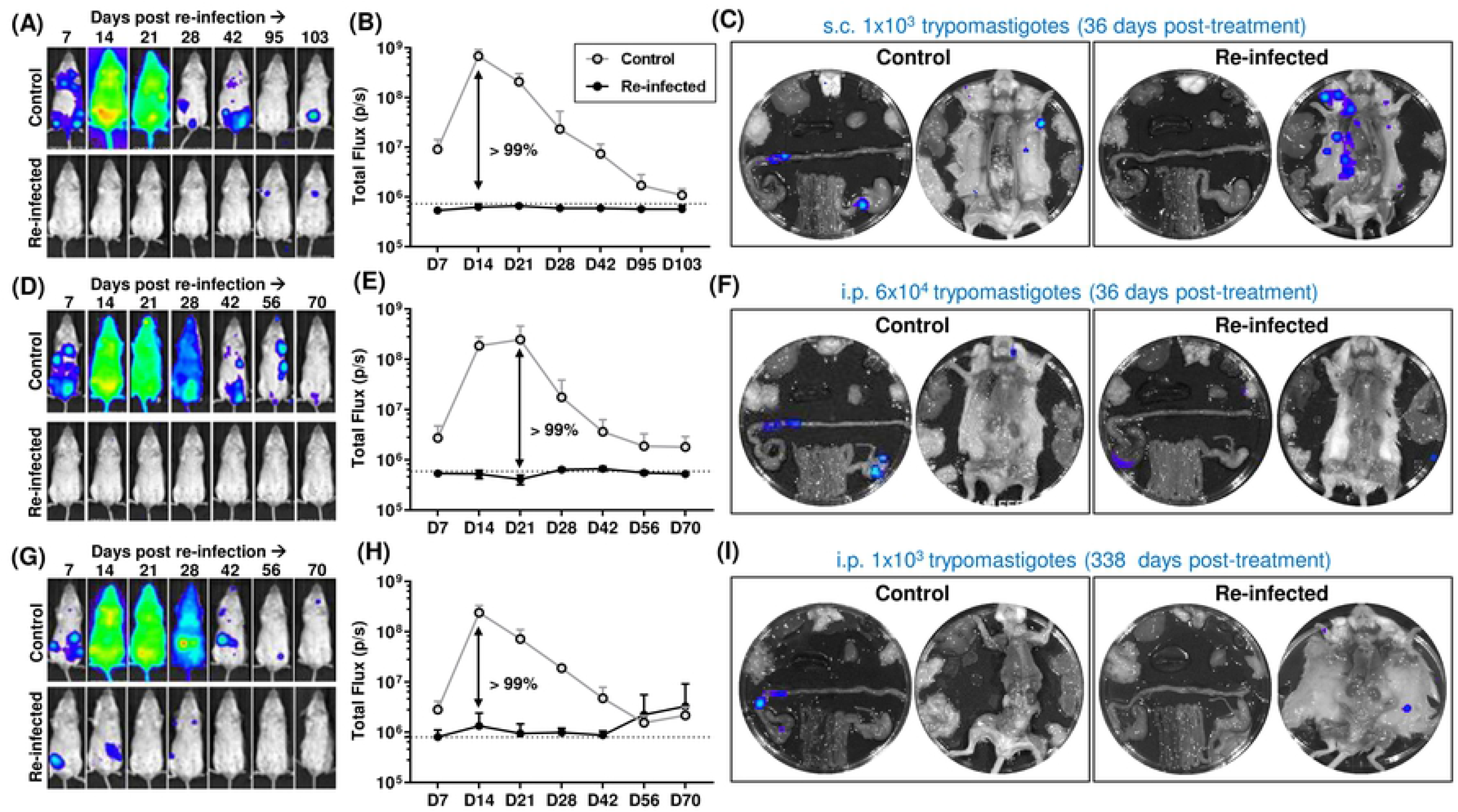
Protection conferred by a benznidazole-cured infection is not dependent on the route of infection, size of the challenge inoculum or the time-period until re-infection. (A) BALB/c mice, infected by the subcutaneous (s.c.) route with 10^3^ bioluminescent trypomastigotes (CL Brener strain), were subjected to curative benznidazole treatment 36 days post-infection. 21 days after the end of treatment, they were re-infected (s.c.). Control mice were also infected by the s.c. route. (B) Total body bioluminescence of drug-cured mice following s.c. re-infection (means ± SD). (C) *Ex vivo* bioluminescence imaging of organs and carcass from a control and the re-infected mouse that was found to be non-protected after immunosuppression (Materials and Methods). (D) BALB/c mice infected i.p. with CL Brener trypomastigotes, were subjected to benznidazole treatment 36 days post-infection. 21 days after the end of treatment, they were re-infected (i.p.) with 6×10^4^ trypomastigotes. (E) Total body bioluminescence of drug-cured mice following re-infection. (F) *Ex vivo* bioluminescence imaging of organs and carcass of a control and the re-infected mouse that was found to be non-protected after immunosuppression. (G) As above, BALB/c mice were infected i.p. and subjected to curative benznidazole treatment. 338 days after the end of treatment, they were re-infected (i.p.) with 10^3^ trypomastigotes. (H) Total body bioluminescence of drug-cured mice following re-infection. The increased mean bioluminescence of the re-infected mice towards the end of the monitoring period was due to an intense bioluminescence focus in one of the two non-protected animals. (I) *Ex vivo* bioluminescence imaging of organs and carcass of a control and a re-infected mouse that was found to be non-protected after immunosuppression. In all cases, the original cohort size was n=6. During the course of the s.c. infection experiment (A-C), one mouse failed to recover from anaesthesia, and was excluded from the analysis.

It has been reported that the capacity of a cured mouse to resist re-infection is dependent on the size of the challenge inoculum (49). To investigate this using the *in vivo* imaging system, mice where curative treatment was initiated 36 days post-infection were challenged i.p. with 6×10^4^ CL Brener trypomastigotes, 60 times the number used previously. The outcome was similar. None of the mice displayed an acute stage parasite burden profile, and only a single mouse (out of 6) was non-protected (Fig 4D, E and F). Therefore, the level of protection conferred by a cured infection is similar when a higher challenge inoculum is used.

We next sought to determine whether protection results from the development of immunological memory, rather than retention of effector cells from the primary infection. BALB/c mice were infected i.p. with CL Brener trypomastigotes, and curative benznidazole treatment was initiated after 36 days. On this occasion, however, the mice were not re-infected until 338 days after the cessation of treatment. Following homologous challenge, the level of protection was comparable to that achieved in previous experiments when mice were reinfected 3 weeks after the last dose of benznidazole. They were able to prevent the onset of parasite proliferation during the acute stage, and 4 out of 6 mice were fully protected (Figs 4G, H, and I).

### Assessing circulating IFN-γ^+^ T cells in mice after primary infection and challenge

We sought to determine if the duration of the primary infection, prior to commencement of curative drug treatment, impacted on the extent of the *T*. cruzi-specific immune response to challenge with homologous parasites. Blood was collected, 2 days prior to re-infection and at regular intervals thereafter, from mice that had been infected for 4, 14 and 36 days (Fig 5A). PBMCs were isolated, re-stimulated with an ASP-2 and TS peptide pool, and the IFN-γ^+^ cell frequencies measured by ELISpot (Materials and Methods). As expected for mice infected with *T. cruzi* (50), the control group receiving their first parasite exposure showed a delayed peptide-specific response, with the frequency of IFN-γ^+^ cells on day 10 not significantly different to pre-infection levels. There were substantial increases by days 25 and 40 (Fig 5B), co-incident with the period when the parasite burden had been controlled and was undergoing major reduction (Figs 3B and C).

**Fig 5.**
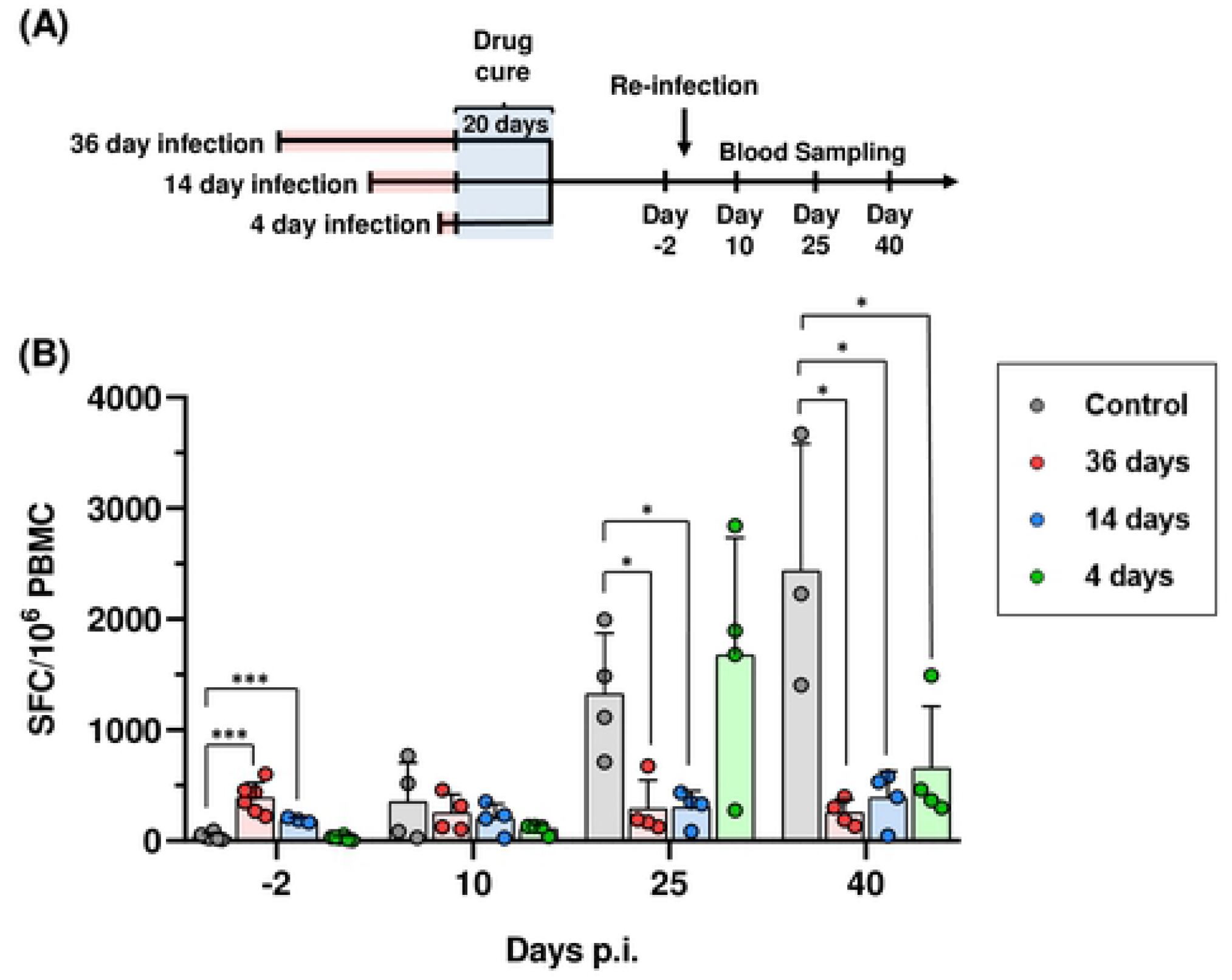
The length of the primary infection affects the level of circulating murine IFN-γ^+^ T cells after challenge. (A) Timeline of experiment. Infected BALB/c mice were subjected to curative benznidazole treatment initiated 36, 14 or 4 days post-infection (as in Figure 3). 21 days after the end of treatment, they were re-infected (i.p., 10^3^ CL Brener trypomastigotes) and blood was collected by venesection on the days indicated. (B) ELISpot analysis. PBMCs were isolated from re-infected mice at pre and post re-infection, as indicated. IFN-γ^+^ PBMCs were quantified after overnight stimulation with a 20-mer peptide pool representing the ASP-2 and TS proteins, as in Figure 1. Data are presented as *ex vivo* IFN-γ SFCs per 10^6^ PBMCs. (***) *p*<0.001; (*) *p*<0.05.

With benznidazole-cured mice, the pre-challenge levels of circulating *T. cruzi-specific* IFN-γ^+^ T cells varied, depending on the duration of the preliminary infection (Fig 5B). Initially, mice that had been infected for 36 days prior to cure displayed higher levels than the control cohort (day-2). However, these levels did not increase significantly following re-infection, although in the case of the day 14 group, there was a slight trend in this direction. In the control group, the robust adaptive response, which controlled the infection, was associated with levels of circulating IFN-γ^+^ T cells that were significantly higher than in any of the benznidazole-cured mice by 40 days post-challenge (Fig 5B). Mice that had been infected for only 4 days prior to drug-cure, displayed IFN-γ^+^ T cell kinetics that were initially more similar to naïve mice receiving their preliminary infection, with very low levels before and 10 days post-challenge, followed by a major increase by day 25. The levels decreased thereafter, in contrast to the control group, where they continued at higher levels until day 40.

We also examined circulating parasite-specific IFN-γ^+^ T cells in mice that had been re-infected 338 days after drug cure (Fig 6A). Prior to re-challenge, the levels were similar to those in non-infected control mice. However, by 10 days post-challenge, there had been a 5-fold increase, in contrast to the delayed peptide-specific response typical of a *T. cruzi* infection (Fig 6B). This difference was not maintained, and on days 25 and 40 after challenge, the level of IFN-γ^+^ T cells was not significantly different from control mice.

**Figure 6.**
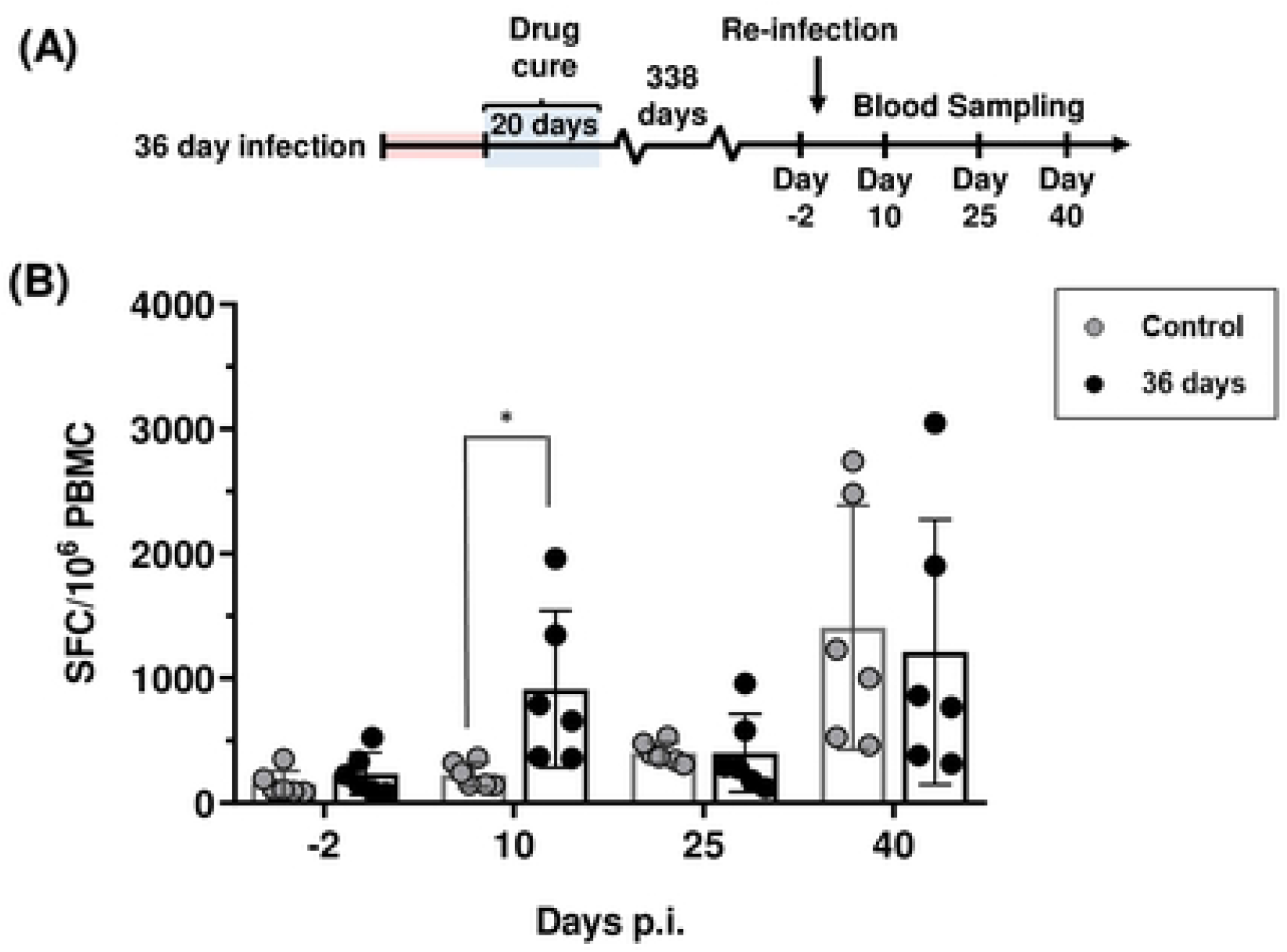
Effect of delaying re-infection on the level of circulating murine IFN-γ^+^ T cells after challenge. (A) Timeline of experiment. Infected BALB/c mice were cured with benznidazole treatment, which was initiated 36 days post-infection, as in Figure 5. They were re-infected 338 days later, and blood samples collected on the days indicated. (B) PBMCs were isolated from re-infected mice at various time points pre and post re-infection. IFN-γ^+^ PBMCs were quantified after overnight stimulation with the 20-mer ASP-2/TS peptide pool. (*) *p*=0.024.

### The capacity of drug-cured infections to confer a cross-strain protective response

*T. cruzi* displays significant genetic diversity, with the natural population subdivided into 6 lineages known as discrete typing units (DTUs), each of which has the ability to infect humans (51). We therefore investigated if the capacity to confer homologous protection is a general feature of *T. cruzi* infections by performing an analogous experiment using JR strain parasites from the genetically distant TcI lineage). In BALB/c mice, infections with this strain are slightly slower to reach the peak of the acute stage, but the bioluminescence profile is otherwise similar to that of the CL Brener strain (DTU VI lineage) (25). Mice were benznidazole-treated 36 days into a JR infection, and re-infected with the same strain 20 days after the end of treatment (Fig 7A). As before, we observed that the cured infection conferred significant protection. None of the mice exhibited a distinct acute stage peak, with the majority remaining close to bioluminescence background levels (Figs 7B and C). However, only a single mouse (out of 6) exhibited sterile protection when assessed by *ex vivo* imaging following immunosuppression (Materials and Methods) (Fig 7D).

**Fig 7.**
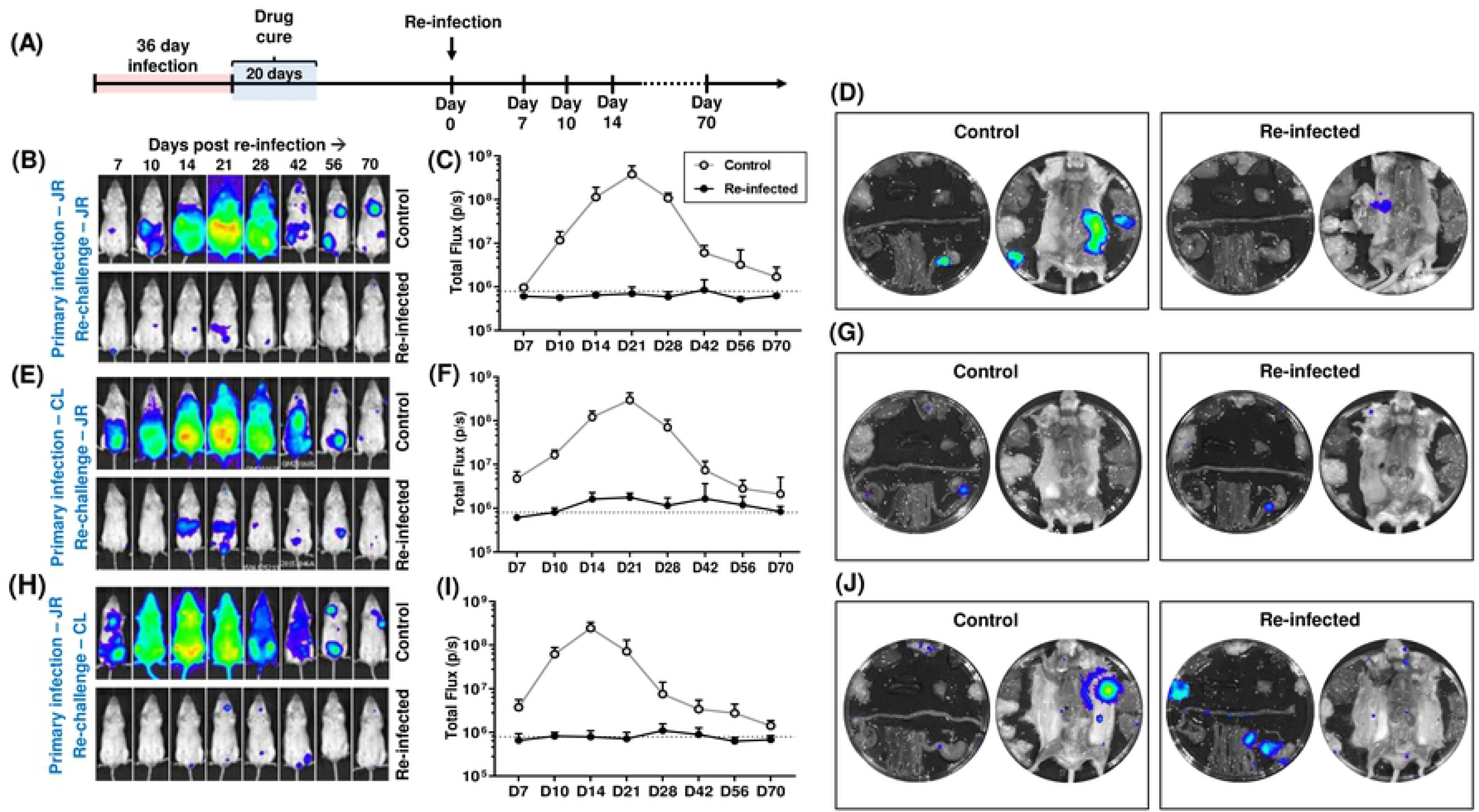
Protection against challenge with a heterologous *T. cruzi* strain. (A) Outline strategy. BALB/c mice were infected i.p. with bioluminescent trypomastigotes (CL Brener or JR strains) and subjected to curative benznidazole treatment initiated 36 days post-infection. 20 days after the end of treatment, they were re-infected as indicated below and monitored for a further 70 days. Bioluminescence-negative mice were then immunosuppressed and assessed by *ex vivo* imaging. (B) Ventral images of a representative drug-cured JR infected mouse (n=6) following re-infection with the homologous JR strain. All images use the same log10-scale heat-map shown in Fig 2. (C) Total body bioluminescence (sum of ventral and dorsal images) of drug-cured JR infected mice re-infected with the JR strain (means ± SD). (D) *Ex vivo* bioluminescence imaging of organs and carcass of a control and re-infected mouse (non-protected). (E) Ventral images of a representative drug-cured CL Brener infected mouse (n=6) following re-infection with the JR strain. (F) Total body bioluminescence of drug-cured CL Brener infected mice re-infected with the JR strain. (G) *Ex vivo* bioluminescence imaging of organs and carcass of a control and re-infected mouse (non-protected). (H) Ventral images of a representative drug-cured JR infected mouse (n=6) following re-infection with the CL Brener strain. (I) Total body bioluminescence of drug-cured JR infected mice re-infected with the CL Brener strain. (J) *Ex vivo* bioluminescence imaging of organs and carcass of a control and re-infected mouse (non-protected).

To assess the scope for vaccine-induced species-wide immunity, we next investigated the effectiveness of protection conferred against a heterologous challenge using the strains CL Brener (TcVI) and JR (TcI). Following the strategy outlined above, BALB/c mice were infected i.p. with 10^3^ bloodstream CL Brener or JR trypomastigotes, benznidazole-treated, and then challenged with the heterologous strain 20 days after the end of the curative therapy. In both experiments, we observed a strong protective response (>99%) (Figs 7E-J), with no distinct acute stage peak and a reduced number of bioluminescent foci in the period corresponding to the transition to the chronic stage. However, in both cases, all mice exposed to cross-strain challenge displayed small but clear parasite foci following re-infection. None exhibited sterile protection when examined by *ex vivo* imaging, with each displaying the type of GI tract infections characteristic of the chronic stage (Figs 7G and J). Therefore, although infection with a heterologous strain can have a major impact on the subsequent parasite burden, it did not confer sterile immunity in any of the mice examined.

## DISCUSSION

Although experimental *T. cruzi* vaccines have been widely shown to reduce the burden of infection in animal models (27–35), there is little unambiguous evidence for sterile protection. Despite this, there have been an increasing number of reports that vaccination could have therapeutic benefits in terms of decreased cardiac pathology (33, 52-54). Therefore, the question as to whether the development of a Chagas disease vaccine might be a practical option for reducing the public health impact of this infection remains unanswered. Detailed assessment has been limited by difficulties in detecting the intermittent low-level parasitemia of chronic stage infections, and in identifying the tissue/organ location of persistent parasites. Here, we demonstrate that highly sensitive bioluminescent imaging can negate some of these issues, and provide novel insights into vaccine efficacy.

Initially, we tested the protective properties of two viral vectors (MVA and ChAdOx1) that had been modified to express an ASP-2/TS fusion gene (Fig 1). The reduction in peak parasite burden (77%, Fig 2C) was in a range similar to that reported for other recombinant *T. cruzi* vaccines (55-57, as examples). Interestingly, vaccination had no impact on the parasite burden once the infection had transitioned to the chronic phase, suggesting that if parasites can survive until this stage of the disease, they are less susceptible to clearance by vaccine-induced immunity. This has not been reported previously. At the experimental end-point, the remaining parasites were restricted predominantly to the GI tract (Fig 2D). In BALB/c and other mice, this location serves as a permissive niche, enabling parasites to persist in an otherwise hostile immune environment (24, 25, 37), although the mechanism(s) for this have yet to be elucidated. To determine if this might limit the utility of a Chagas disease vaccine, we therefore investigated the protective effect of benznidazole-cured infections, on the basis that these should provide optimal levels of immunity. Curative treatment has been associated with the development of a stable anti-*T. cruzi* CD8^+^ T cell population (22).

Drug-cure was initiated after infection for 36 days (a time-point when the adaptive CD8^+^ T cell response is controlling the infection), 14 days (the peak of the acute stage), and 4 days (just short of one complete round of the intracellular replication cycle that leads to differentiation and parasite egress) (Fig 3). Mice infected for 36 days prior to benznidazole treatment were highly protected from challenge (Fig 3B), with complete absence of a typical acute stage peak. However, sterile protection was only achieved in half of the re-infected mice (Table 1). Therefore, although drug-cured infections can generate a highly effective immune response, that prevents a second acute phase, parasites that evade this initial encounter seem to be refractory to immune-mediated elimination and are able to persist long term. This outcome was not significantly influenced by the size of the challenge inoculum or the route of infection (Fig 4). Initiating treatment after 14 days was also able to prevent a detectable acute stage peak in bioluminescence when the mice were challenged, although there was a slight reduction in the level of sterile protection (Fig 3B, Table 1). In contrast, when drug-cure was initiated after 4 days, a pronounced post-challenge acute stage peak could be observed, but even then, the parasite burden was 85% lower than in a naive infection. Therefore, 14 days exposure to an untreated infection is sufficient for the induction of a robust immune response, whereas with 4 days, the response appears to be less developed, although still sufficient to have a significant impact on the parasite burden.

A delayed onset of the CD8^+^ T cell response is a characteristic feature of Chagas disease (58), with the first round of intracellular infections passing largely undetected by the immune system. Upon invasion of mammalian cells, parasites rapidly escape from the phagolysosome, there is down-modulation of the host cell immunoproteasome (59), and minimal activation of the host-pattern recognition receptors. Induction of effective innate immunity requires a full cycle of parasite replication, host cell lysis, and the release of trypomastigotes into the extracellular milieu, a process that takes at least 4-5 days. This is followed by the production of pathogen-associated and damage-associated molecular patterns that promote innate immune responses, allowing parasitized cells to flag up their infected status by MHC class I antigen presentation. Full development of the CD8^+^ T cell response to *T. cruzi* infections takes around 3 weeks (58). In mice where curative treatment was initiated 14 or 36 days postinfection, circulating parasite peptide-specific IFN-γ^+^ T cells were readily detectable prior to challenge (Fig 5B), and were associated with protection against the development of a second acute phase profile. In many cases, this response was sufficient to promote complete elimination of the secondary infection. Experimental challenge did not lead to a significant increase in the level of IFN-γ^+^ T cells, suggesting that the pre-existing effector population was able to contain the secondary infection without further induction. Even when preliminary infection did not confer sterile protection, there was no further enhancement of the peptide-specific response. Therefore, if parasites in the challenge inoculation can avoid early elimination, and are able to establish a long term chronic infection, it appears that they survive in an environment or state that does not trigger additional T cell activation.

In mice where curative treatment was initiated 4 days into the primary infection, the level of *T. cruzi*-specific IFN-γ^+^ T cells prior to re-infection was negligible, and the kinetics of the response induced over the first 25 days of the challenge infection was similar to that in the controls (Fig 5). Despite this, there was an 85% reduction in the parasite burden at the peak of the re-infection. Therefore, the induced partially protective effect in these mice is conferred either by an extremely low level of circulating parasite-specific IFN-γ^+^ T cells, or by other factors that operate to moderate the infection. In mice challenged almost a year after curative treatment of the primary infection, the level of protection was similar to mice in which the gap between the end of treatment and challenge was only ~20 days. However, unlike these mice, re-infection after almost a year was accompanied by induction of peptide-specific T cells, with kinetics that were more rapid than in the naive control cohort (Fig 6). This evidence for a memory response suggests that vaccine-mediated long term protection against fulminant *T. cruzi* infection may be a feasible goal. Furthermore, if these results can be extrapolated to humans, it would imply that patients who have undergone curative drug treatment should have the added benefit of a high level of long term protection against re-infection.

We also investigated the extent to which benznidazole-cured infections could provide cross-strain protection. *T. cruzi* is highly diverse, with six major genetic lineages that display considerable geographic overlap. Taxonomy is further complicated by the widespread existence of hybrid strains (60). Mice initially infected with the *T. cruzi* CL Brener (TcVI) were challenged with the JR strain (TcI), and vice-versa. Although, suppression of the parasite burden was similar to that in a homologous challenge (>99%), sterile protection was not achieved (Fig 7), with surviving parasites persisting at very low levels. We have suggested a model for chronic Chagas disease (37) in which the gut (and perhaps other tissues, such as the skin or skeletal muscle) acts as an immunologically tolerant reservoir for *T. cruzi* persistence, with periodic trafficking to other sites, where the parasites are then destroyed rapidly by immune effector mechanisms. In the heart, this can lead to cumulative collateral damage that ultimately gives rise to cardiac pathology (61).

We propose that when parasites establish infections in the GI tract, or other permissive sites, they become refractory to elimination by the vigorous adaptive responses induced by drug-cured infections. Thus, the effectiveness of a Chagas disease vaccine could depend on the efficiency with which the primed immune system prevents *T. cruzi* from reaching the relative safety of these sites of persistence, and its ability to maintain this response over time against a wide range of strains. The results presented here, and elsewhere (31–34, 44, 45, 52–57), highlight the possibility that current subunit/DNA vaccines may be unable to fulfil these requirements, although their ability to prevent lethal outcomes (31–34, 44, 45, 53, 56) and provide therapeutic benefits (33, 52–55, 57) merits further research. As we have shown here however, the long term protection conferred by live infection, followed by drug-mediated cure, suggests that the use of genetically attenuated parasite strains may be the best approach to achieving an effective vaccine.

## ACKNOWLEDGEMENTS

We thank Helena Helmby and Julius Hafalla (LSHTM) for advice and guidance.

## SUPPLEMENTARY INFORMATION

**Supplementary Figure 1. Assessing the curative ability of benznidazole**. (A, B) *In vivo* imaging of BALB/c mice infected with CL Brener (A) and (JR) strains of *T. cruzi*. Treatment with benznidazole, 100 mg kg^-1^ once daily by the oral route for 20 days, was initiated 36 days post-infection. Following cessation of treatment, mice were immunosuppressed with 3 doses of 200 mg kg^-1^ cyclophosphamide (Materials and Methods). All images use the same log_10_-scale heat-map with minimum and maximum radiance values indicated. (C and D) Total body bioluminescence (sum of ventral and dorsal images) of CL Brener (C) and JR (D) infected mice. Dashed lines indicate background bioluminescence. All images use the same log_10_-scale heat-map with minimum and maximum radiance values indicated. (E and F) *Ex vivo* bioluminescence imaging of organs and carcasses from CL Brener (E) and JR (F) infected mice at the experimental end-point. A minor bioluminescent focus was observed in the adipose tissue of mouse 1 (JR infection). Mouse 2 (CL Brener infection) was euthanised prior to day 89, due to weight loss during immunosuppressive treatment. It was negative by both *in vivo* and *ex vivo* imaging.

**Supplementary Figure 2. Benznidazole-cured infections confer significant protection against re-challenge with the *T. cruzi* CL Brener strain.** (A) Timeline. BALB/c mice infected i.p. with 10^3^ trypomastigotes (CL Brener strain) were subjected to curative benznidazole treatment initiated 36 days post-infection. 23 days after the end of treatment, they were re-infected i.p. After a further 75 days, the mice were immunosuppressed using cyclophosphamide (red stars) and assessed by *ex vivo* imaging. (B) Ventral and dorsal bioluminescence images from a cohort of 6 mice. The days post re-infection are indicated (left). All images use the same log10 scale heat-map with minimum and maximum radiance values indicated. (C) *Ex vivo* bioluminescence imaging of organs and carcasses harvested at the experimental end-point. (D) Total body bioluminescence (sum of ventral and dorsal images) of drug-cured mice following re-infection (means ± SD) derived by *in vivo* imaging. Mice 3 and 4 were designated as non-protected on the basis of *in vivo* and/or *ex vivo* imaging.

